# Tendon response to matrix unloading is determined by the patho-physiological niche

**DOI:** 10.1101/620534

**Authors:** Stefania L. Wunderli, Ulrich Blache, Agnese Beretta Piccoli, Barbara Niederöst, Claude N. Holenstein, Fabian Passini, Unai Silván, Louise Bundgaard, Ulrich auf dem Keller, Jess G. Snedeker

## Abstract

Aberrant matrix turnover with elevated matrix proteolysis is a hallmark of tendon pathology. While tendon disease mechanisms remain obscure, mechanical cues are central regulators. Unloading of tendon explants in standard culture conditions provokes rapid cell-mediated tissue breakdown. Here we show that biological response to tissue unloading depends on the mimicked physiological context. Our experiments reveal that explanted tendon tissues remain functionally stable in a simulated avascular niche of low temperature and oxygen, regardless of the presence of serum. This hyperthermic and hyperoxic niche-dependent catabolic switch was shown by whole transcriptome analysis (RNA-seq) to be a strong pathological driver of an immune-modulatory phenotype, with a stress response to reactive oxygen species (ROS) and associated activation of catabolic extracellular matrix proteolysis that involved lysosomal activation and transcription of a range of proteolytic enzymes. Secretomic and degradomic analysis through terminal amine isotopic labeling of substrates (TAILS) confirmed that proteolytic activity in unloaded tissues was strongly niche dependent. Through targeted pharmacological inhibition we isolated ROS mediated oxidative stress as a major checkpoint for matrix proteolysis. We conclude from these data that the tendon stromal compartment responds to traumatic mechanical unloading in a manner that is highly dependent on the extrinsic niche, with oxidative stress response gating the proteolytic breakdown of the functional collagen backbone.

## Introduction

Tendon is a highly specialized connective tissue that consists largely of sparsely distributed stromal tendon fibroblasts in a collagen-rich extracellular matrix (ECM). The fact that tendons bear physiologically extreme mechanical stresses is reflected by its tightly packed and highly structured ECM. The load bearing tendon core is comprised by hierarchically organized type-I collagen fibrils, fibres and fascicles (or fibre bundles) [1], with fascicles representing the basic functional load-bearing unit of stromal tissue [2, 3]. Physiological mechanical signals are understood to regulate the lifelong adaption of the tissue to individual functional demands [4–8], with a narrow gap between beneficial and detrimental mechanical loads [9–12]. This threshold is often exceeded, with tendon disorders accounting for 30 – 50% of all musculoskeletal clinical complaints associated with pain [13].

Our basic knowledge of the molecular and cellular mechanisms behind tendon physiology and pathology is limited [14–16], yet damage and inadequate repair are considered to be central to tendon disease onset and progression [17–19]. The stimuli that trigger tissue healing responses remain unclear, potentially being a direct effect of cellular activation by overloading or by unloading after isolated fibre rupture with subsequently altered cell-matrix interaction [20]. However, the mechanisms of overloading and underloading may not be mutually exclusive, with tissue damage resulting in localized regions of both load types. Regardless of the mechanism of damage activation, aberrant healing processes typically result in a relatively disorganized and mechanically inferior repair tissue [21–23]. Tendinopathy severity is graded by its histopathological score, characterized by the extent of disorganized collagen matrix, abnormal cell shape, presence of inflammation, increased cellularity, vascularization and neuronal ingrowth [24–28]. These hallmarks of disease intimately involve tissue turnover and proteolytic breakdown of the ECM [29–32]. Although pathological matrix turnover is central to tendinopathy, very little is known about either the initiation or regulation of tissue catabolism in the tendon core [33–36].

The healthy tendon stromal niche represents a low-oxygen and low-temperature tissue that is minimally vascularized. Tendon fibroblasts reside in a state of relative metabolic quiescence compared to stromal cells in other tissues [37]. Tendon repair thus likely involves closely coordinated interaction between the hypovascularized intrinsic tendon core and the peritendinous synovial compartment of the tissue that connects to the vascular, immune and nervous systems [38]. Although a clear picture is only emerging, we speculate that mechanical sensors within the intrinsic tendon compartment recruit structures of the extrinsic compartment resulting in neo-vascularization, neuronal ingrowth, immune response and inflammation (Figure 1A) [38]. While patho-histological investigations of symptomatic tendon phenotypes clearly demonstrate the presence of blood vessels within the tendon proper, it remains unclear, which signalling-cascades are initiated in the intrinsic compartment by external factors [39–42].

**Figure 1:**
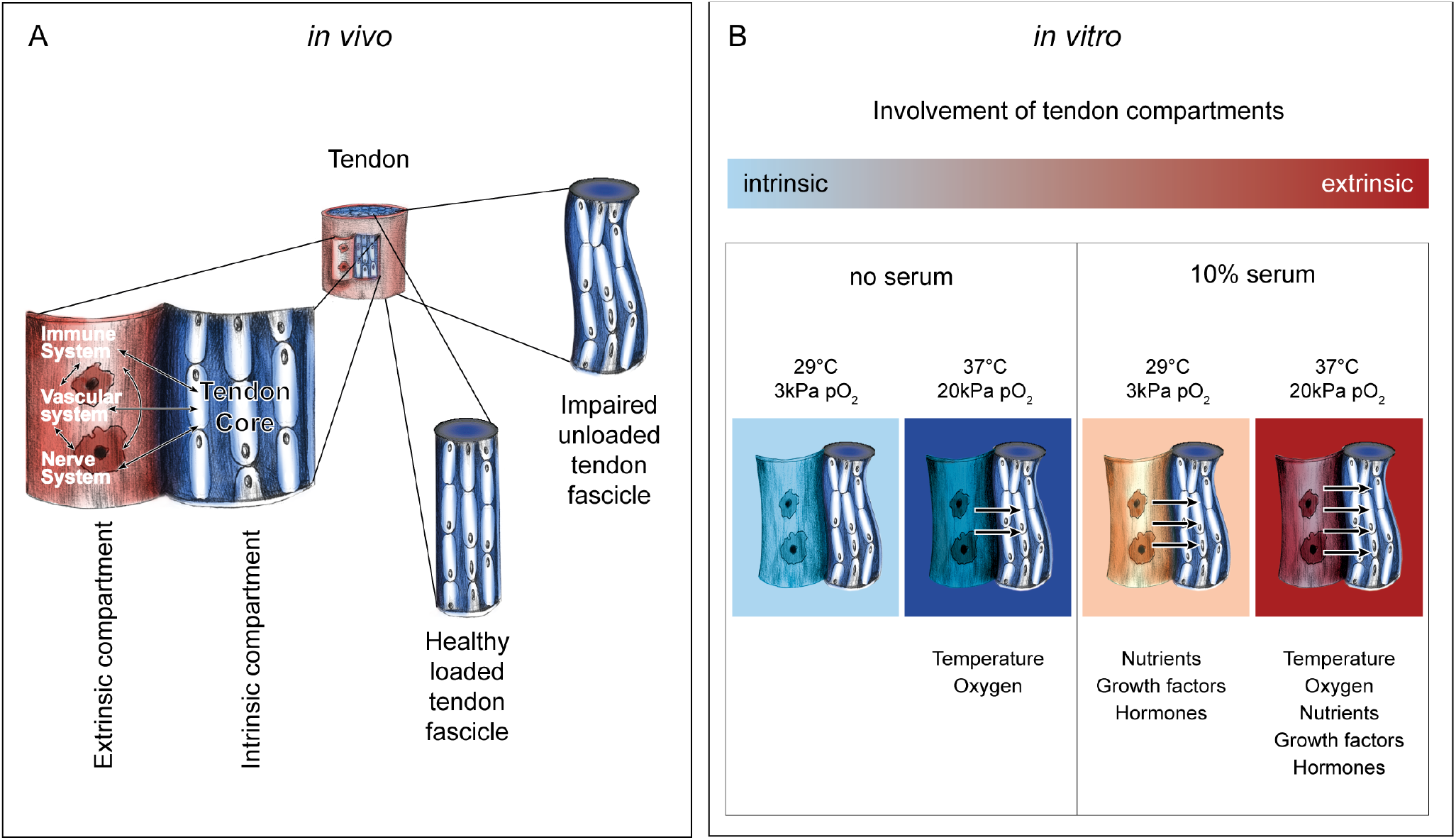
Tendon *in vivo* and *in vitro* model system. **(A)** Tendon is a hierarchically structured tissue. It mainly consists of densely packed collagen type I matrix and tendon cells (tenocytes) that are responsible for tissue maintenance. The tendon fascicle is the basic functional unit of the tendon (tendon core) and represents the intrinsic tendon compartment. The extrinsic compartment comprises the wrapping layer with synovium-like character that connects immune, vascular and nervous system. Wrapping layers are present at several hierarchical levels and are named endotendon, epitenon and paratenon (from lowest to highest level). It is thought that intrinsic and extrinsic compartments interact with each other at all levels to ensure proper tissue maintenance during homeostasis or repair. In a healthy homeostatic tissue tendon fascicles are loaded, while unloading of tendon fascicles may occur due to overloading and subsequent rupture of sub-tissue structures or upon immobilization. **(B)** In our *ex vivo* model the impaired tendon core is represented by an unloaded murine tail tendon fascicle. By varying temperature, oxygen and serum content of the culture conditions we mimicked the degree of involvement of the extrinsic compartment on the cell-driven healing process.

In this study we investigated how deviations from a low-oxygen, low-temperature tissue niche affect resident stromal cells, specifically hypothesizing that the tissue niche is an important regulator of cellular response to matrix damage. We established a system in which explanted murine tail tendon fascicles served as a model of an independent unit of the intrinsic compartment that had been unloaded by mechanical tissue damage. Unloaded tendon fascicles were exposed to different niches of varying temperature, oxygenation and nutrient supply to mimic pathological and vascularized states of different severity (Figure 1B). We compared mechanical properties, cell viability, metabolic activity, tissue morphology, whole transcriptome, and protein degradomes and secretomes. Hereby we identified mechanisms of extrinsic tendon compartment regulation on the response of intrinsic tendon cells towards tissue homeostasis or pathologic tissue degeneration by activating an immune-related cellular response, production of reactive oxygen species and downstream activation of proteolytic enzymes.

## Methods

### *Ex vivo* tissue culture

Tendon fascicles were extracted from tails of freshly euthanized 12-13 weeks old wild-type C57BL6/J male mice. Tendon fascicles were gently isolated from each tail using a surgical clamp and distributed among the control and treatment groups in a non-biased fashion. The explanted fascicles were cultivated in absence of mechanical load and examined at designated time-points (6, 12, 18, 24 days). Freshly isolated fascicles served as a positive tissue control (day 0). Devitalized fascicles were generated by three freeze-thaw cycles before further use and served as negative control. Fascicles were cultured in serum-free medium (high glucose Dulbecco’s Modified Eagle’s Medium (DMEM), 1% N2-supplement (v/v), 1% penicillin-streptomycin (v/v), 200μM L-ascorbic acid phosphate magnesium salt n-hydrate) or serum-supplemented medium (high glucose DMEM, 10% fetal bovine serum (v/v), 1% penicillin-streptomycin (v/v), 1% non-essential amino acids (v/v), 200μM L-ascorbic acid phosphate magnesium salt n-hydrate). Fascicles were maintained in a humidified atmosphere containing 5% CO_2_ at either low oxygen partial pressure (*pO_2_* = 3kPa) and low temperature (29 °C) or high oxygen partial pressure (*pO_2_* = 20kPa) and temperature (37°C). The fascicles in serum-supplemented medium were additionally cultured in intermediate conditions at high oxygen and low temperature or vice versa. Medium was changed every sixth day. All compounds of the cell culture media were purchased from Sigma Aldrich, except for the ascorbic acid (Wako Chemicals) and the N2-supplement (Gibco). The Cantonal Veterinary office of Zürich ethically approved all animal experiments (permit number ZH265/14) and a total number of 40 mice was used.

Physiological tail tendon temperature of healthy mice was estimated by infrared-measurements of the cutaneous tail temperature in 40 animals (Infrared thermometer 153-IRB, Bioseb).

### Biomechanical testing and analysis

Force-displacement data were recorded on a custom-made uniaxial stretch device as previously described [43]. In brief, fascicles were clamped, pre-loaded to 0.015N (corresponding to 0% strain) and preconditioned to 1% strain with five stretch cycles (n=6, independent samples from different mice). The tangential elastic-modulus was calculated in the linear part of the stress-strain curve (0.5 – 1% strain) of a single additional stretch cycle to 1% by fitting a linear slope (Matlab R2016a, Version 9.0.0.341360). The pre-load was reapplied after every cycle. Nominal stress was calculated based on the initial cross-sectional area. Cross-sectional area was determined from the diameter assuming a round specimen shape. Diameter was measured using automated evaluation of microscopic images (Matlab R2016a, Version 9.0.0.341360), which were recorded before the treatment at three different locations along the fascicle with a 20-fold magnification (Motic AE2000, Plan LWD 20x, WD 4.7).

### Cell viability and ATP measurements

Dead cells in the fascicles were stained with Ethidium Homodimer-1 (2μM, AS 83208, Anaspec) previous to 10% formalin fixation (HT5011, Sigma Aldrich). Subsequently, cell nuclei were stained with NucBlue reagent (1 drop/ml = ca. 10μl, R37606, Thermo Fischer, Switzerland) and embedded in a 50% Glycerol:PBS solution. Z-stack fluorescent images were obtained in triplicates by using a spinning disc confocal microscope (iMic, FEI) equipped with a Hamamatsu Flash 4.0 sCMOS camera and a SOLE-6 Quad laser (Omicron) and using a 20x objective (Olympus, UPLSAPO, N.A. 0.75). Total number of imaged cells, percentage live cells (total cell number minus dead cells) and fascicle volume (n=6, independent samples from different mice) were assessed by means of a custom microscopic image analysis tool (MATLAB, R2016a 9.00.34160). Number of cells was determined in a measurement depth of 60μm (from surface) and the volume of the circular segment was calculated by measuring length and diameter of total specimen z-projection.

ATP was assessed by using a luminescence based ATP detection assay according to the manufacturer’s instructions (CellTiterGlo3D, G9681, Promega). ATP standards were prepared from a 10mM stock solution (P1132, Promega). After ATP measurement the 20mm long fascicle parts (n=4 in duplicate measurements, two fascicles per animal) were frozen at −80°C and subsequently digested at 56°C overnight by Proteinase K (10mM in Tris-HCl, 19133, Qiagen). To account for potentially differential cell proliferation rate between treatments all the ATP quantifications were normalized to DNA content in the same samples. The quantity of DNA in each digest was measured by fluorescent signal detection using the Quant-iT™ dsDNA High-Sensitivity Assay Kit (Q33232, Thermo Fisher) according to the manufacturer’s protocol. Whole fascicles were cut to equally sized parts (20mm), while one half was processed and measured directly after isolation (day 0) and the other half after the treatment (day 6 and 12). Lumiescence and fluorescence were both recorded on a microplate reader (SpectraMAX Gemini XS).

### Second harmonic generation imaging and transmission electron microscopy of tendon fascicles

Collagen was visualized by second harmonic generation (SHG) imaging of whole mount tissues on a multi-photon and confocal fluorescence microscope (Leica TCS SP8) equipped with photon-counting hybrid detectors and an ultrafast near infrared laser (Insight DS+ Dual, Spectra Physics) using 25x magnification (HC IRAPO L 25x, NA 1.0), an excitation wavelength of 880nm and a SHG-compatible fluorescence filter cube (440/20 & 483/32 with 455 SP). Cell nuclei were imaged from the same samples. All the samples were recorded at the same laser power and gain control with a z-step size of 0.3μm. After image acquisition 70 stacks were z-projected using ImageJ (Fiji, NIH Version 1.0).

For transmission electron microscopy whole mouse tail tendon fascicles were fixed in 2.5% glutaraldehyde (G5882, Sigma-Aldrich) in 0.1M Cacodylat buffer (pH 7.2) for 20 minutes. To visualize the collagen fibrils, samples were contrasted with 1% uranyl acetate in water for 1 h. All samples were then dehydrated with graduated concentration of ethanol and propylene oxide. Specimens were infiltrated and embedded in Epon (polymerized at 60 °C for 48 h). Ultra-thin longitudinal and cross sections were cut with a diamond knife and mounted on uncoated grids. Samples were then imaged with a transmission electron microscope (TEM-FEI Tecnai G2 Spirit).

### RNA sequencing

For each experimental group 16-20 fascicles were pooled after 6 days of culture, snap frozen in liquid nitrogen and pulverized in QIAzol lysis reagent (#79306, Qiagen) by cryogenic grinding (FreezerMill 6870, SPEX^TM^ SamplePrep). Tissue lysates were mixed with 1-bromo-3-chloropropane (B9673, Sigma-Aldrich) at a ratio of 1:4 and the RNA containing aqueous/organic phase was separated using 5Prime PhaseLock Gel Heavy (#2302830, LabForce). RNA was extracted by using the RNeasy micro Kit (74004, Quiagen). The quantity and quality of the isolated RNA (n = 3, independent pools of fascicles from different mice) was determined with a Qubit® (1.0) Fluorometer (Life Technologies, California, USA) and a Bioanalyzer 2100 (Agilent, Waldbronn, Germany). The TruSeq Stranded mRNA Sample Prep Kit (Illumina, Inc, California, USA) was used in the succeeding steps. Briefly, total RNA samples (100 ng) were poly-A selected and then reverse-transcribed into double-stranded cDNA with Actinomycin added during first-strand synthesis. The cDNA sample was fragmented, end-repaired and adenylated before ligation of TruSeq adapters. The adapters contain the index for multiplexing. Fragments containing TruSeq adapters on both ends were selectively enriched with PCR. The quality and quantity of the enriched libraries were validated using Qubit® (1.0) Fluorometer and the Bioanalyzer 2100 (Agilent, Waldbronn, Germany). The product is a smear with an average fragment size of approximately 360 bp. The libraries were normalized to 10nM in Tris-Cl 10 mM, pH8.5 with 0.1% Tween 20. The TruSeq SR Cluster Kit v4-cBot-HS (Illumina, Inc, California, USA) was used for cluster generation using 8pM of pooled normalized libraries on the cBOT. Sequencing were performed on the Illumina HiSeq 4000 single end 125 bp using the TruSeq SBS Kit v4-HS (Illumina, Inc, California, USA).

### RNA sequencing data analysis

Bioinformatic analysis was performed using the R package ezRun^1^ within the data analysis framework SUSHI [44]. In details, the raw reads were quality checked using Fastqc^2^ and FastQ Screen^3^. Quality controlled reads (adaptor trimmed, first 5 and last 6 bases hard trimmed, minimum average quality Q10, minimum tail quality Q10, minimum read length 20nt) were aligned to the reference genome (Ensembl GRCm38.p5) using the STAR aligner [45]. Expression counts were computed using featureCounts in the Bioconductor package Subread [46]. Differential expression analysis was performed using the DESeq2 package [47], where raw read counts were normalized using the quantile method, and differential expression was computed using the quasi-likelihood (QL) F-test^4^. Gene ontology (GO) enrichment analysis was performed using Bioconductor packages goseq [48] and GOStats [49]. Quality checkpoints [50], such as quality control of the alignment and count results, were implemented in ezRun and applied through out the analysis workflow to ensure correct data interpretation. Differentially expressed genes were considered at a p-value <0.05 and a log2ratio > 1. Pathway analyses were performed using MetaCore database via GeneGo tool from Thomson Reuters^5^ and the Kyoto Encyclopedia of Genes and Genomes (KEGG) database.

### Detection of reactive oxygen species

To quantify the production of reactive oxygen species (ROS) tendon fascicles (n=6, independent samples from different mice) were stained with 2’,7’-Dichlorofluorescin diacetate (DCFH-DA, D6883, Sigma Aldrich). Control fascicles and cultured fascicles were incubated for 30 minutes with 50μM DCFH-DA in PBS at either 29°C, 3kPa pO_2_ or 37°C, 20kPa pO_2_ at day 0 and day 6 according to the previous culture. Immediately after incubation, fluorescence was recorded by acquiring z-stacks at steps of 1 μm through the whole fascicles depth at an emission wavelength of 488nm on the spinning disc confocal microscope setup as described above using a 10x objective (Olympus, UPLSAPO, N.A. 0.4). Quantification of fluorescent signal reflected from the tissue was evaluated by measuring cumulative mean intensity of z-projections using ImageJ (Fiji, NIH Version 1.0). Images were taken in triplicates.

### Quantitative real-time polymerase chain reaction (qRT-PCR)

RNA was extracted as described above but using the PureLink™ RNA Micro Scale Kit (#12183016, Invitrogen). cDNA was prepared by reverse transcribing 99ng total RNA in a volume of 44μl using the High-Capacity RNA-to-cDNA™ Kit (#4387406, Applied Biosystems). qRT-PCR was carried out using 2μl cDNA, 5μl TaqMan Universal Master Mix, 0.5μl TaqMan Primer (Applied Biosystems) and 2.5μl RNase free water. Samples were amplified in 40 cycles on a StepOnePlus™ thermocycler (Applied Biosystems) and reactions were run in technical duplicates. Following primers were used: Mm00442991_m1 (*MMP9*), Mm01168399_m1 (*MMP10*), Mm00439491_m1 (*MMP13*), Mm01310506_m1 (*Ctsb*), Mm00515580_m1 (*Ctsc*), Mm00515586_m1 (*Ctsd*), Mm01255859_m1 (*Ctss*), Mm00517697_m1 (*Ctsz*). Relative gene expression was calculated by the comparative Ct method [51] by normalizing the data to the stable housekeeping gene Mm0129059_m1 (*Anxa5)* and to freshly isolated samples.

### Inhibition experiments

Tendon fascicles were cultured under degrading conditions (serum, 37°C and 20kPa pO_2_) for 12 days. From the beginning of the incubation time medium was supplemented with the ROS inhibitor Tempol (4-hydroxy-2,2,6,6-tetramethylpiperidin-1-oxyl)) (1mM, 176141-1G, Sigma), the collagenase inhibitor Ilomastat ((R)-*N*′-Hydroxy-*N*-[(S)-2-indol-3-yl-1-(methylcarbamoyl)ethyl]-2-isobutylsuccinamide) (25μM, CC1010, Merck Millipore) or a combination of both. Medium with DMSO (#472301, Sigma) was used as a control. Culture medium was changed after 6 days. Tissue stiffness was measured at day 0 and day 12 and negative control fascicles were devitalized as described above.

### Mass spectrometry-based proteomics

Terminal amine isotope labeling of substrates (TAILS) analysis, whereby protein N-termini were enriched from 12 day cultures of mechanically intact and impaired fascicles, was performed according to the previously described protocol [52] except for using 10plex Tandem mass Tag™ labelling (TMT, #90110, Thermo Fisher) and adding the trypsin at a ratio of 1:20 (trypsin/protein ratio). Prior to protein denaturation and labelling, proteins of samples were extracted by pressure cycling technology (60 cycles of 45’000psi for 20s, atmospheric pressure for 10s, 33°C) from ca. 4mg tissue in a buffer containing 4M Guanidine hydrochloride (Sigma-Aldrich), 250mM HEPES (Sigma-Aldrich (pH 7.8) and protease inhibitor cocktail (cOomplete EDTA-free, Roche) using a Barocycler 2320ETX (Pressure Biosciences Inc.) (Figure 6A).

Secretomes of 18-22 equally sized and 12-days-cultured fascicles were collected for each of the two conditions in 2ml serum-free and phenol red-free DMEM (#31053028, Gibco supplemented with 1% penicillin-streptomycin (v/v), 200μM L-ascorbic acid phosphate magnesium salt n-hydrate and 1% GlutaMAX (v/v), A1286001, Gibco) during an exposure time of 48 hours. Fascicles were thoroughly washed with PBS before secretome collection. Secretomes were supplemented with protease inhibitor cocktail (cOomplete EDTA-free, Roche) and concentrated ca. 40 times by centrifugal filtration (3K Amicon Ultra, Merck Millipore, 14’000g, 4°C) and buffer was exchanged to 4M Guanidine hydrochloride in 50mM HEPES (pH 7.8). Subsequently, secretomes were reduced in tris(2-carboxyethyl)phosphine (200mM, Sigma-Aldrich) and alkylated in chloroacetamide (400mM, Sigma-Aldrich) for 30 minutes at room temperature and then digested by LysC (Lysyl endopeptidase, #125-05061, Wako, ratio LysC/protein 1:100) and trypsin (Trypsin Gold, V5280, Promega, ratio trypsin/protein 1:20) overnight. Desalting was performed according to the TAILS protocol [52].

PreTAILS, TAILS and secretome peptide mixtures were loaded onto an PepMap100 C18 precolumn (75 μm x 2 cm, 3μm, 100Å, nanoViper, Thermo Scientific) and separated on a PepMap RSLC C18 analytical column (75 μm x 50 cm, 2μm, 100Å, nanoViper, Thermo Scientific) using an EASY-nLC™ 1000 liquid chromatography system (Thermo Scientific) coupled in line with a Q Exactive mass spectrometer (Thermo Scientific). Separation of 1 μg of peptide mixture was achieved by running a constant flow rate of 250 nL/min in 0.1% formic acid/99.9% water and a 140 min gradient from 10% to 95% (23% for 85min, 38% for 30min, 60% for 10min, 95% for 15min) elution buffer (80% acetonitrile, 0.1% formic acid, 19.9% water). A data-dependent acquisition (DDA) method operated under Xcalibur 3.1.66.10 was applied to record the spectra. The full scan MS spectra (300-1750 m/z) were acquired with a resolution of 70,000 after accumulation to a target value of 3e6. Higher-energy collisional dissociation (HCD) MS/MS spectra with fixed first mass of 120 m/z were recorded from the 10 most intense signals with a resolution of 35,000 applying an automatic gain control of 1e6 and a maximum injection time of 120ms. Precursors were isolated using a window of 1.6 m/z and fragmented with a normalized collision energy (NCE) of 28%. Precursor ions with unassigned or single charge state were rejected and precursor masses already selected for MS/MS were excluded for further selection for 60s. Settings for the scan of the secretome peptide mixture were the same except for using a resolution of 17,500 for recording the MS/MS spectra, a maximum injection time of 60ms, and a NCE of 25%.

### Proteomics data analysis

Raw data files were analysed using Proteome Discoverer^TM^ 2.3 software (PD) searching the peak lists against a database collated in the UniProt reference proteome for the species *Mus musculus* (ID: 10090, entries: 17’005). Search parameters for the TAILS experiment were set according to the previously published protocol [52], treating preTAILS and TAILS samples as fractions. For protein-level analysis, quantifiable internal fully tryptic peptides and mature protein N termini were annotated with help of TAILS-Annotator [53], extracted from PD peptide groups and their normalized, scaled abundances were summed up per corresponding protein. The resulting list was curated for protein FDR<0.01 based on PD protein analysis and proteins with significantly differential abundance between conditions determined applying limma moderated t-statistics with Bonferroni-Hochberg multiple testing correction in CARMAweb [54]. Similarly, quantifiable N-terminal peptides were extracted from the PD analysis, annotated using TopFIND [55] and analysed for statistically differential abundance using CARMAweb applying the same parameters as for proteins based on scaled abundances summed up per stripped peptide sequence. Volcano plots were generated using GraphPad Prism (GraphPad Software 8.0).

Raw data files of the secretome experiments were processed by PD using the label-free quantification algorithm Minora Feature Detector node and proteins tested for differential abundance between conditions applying limma moderated t-statistics using CARMAweb [54] on normalized scaled abundance values exported from PD.

### Statistics

RStudio (Version 1.1.423, 2016, Boston, MA) was used for statistical tests and graph illustrations [56]. Mean values of elastic moduli were compared by one-way analysis of variance (ANOVA) followed by contrast analysis. Additionally, multiple linear regression analysis was used to assess the linear relationship between culture time-points, temperature, oxygen and serum as predictors for the response variable elastic modulus. Temperature, oxygen and serum were coded as 0 = 29°C / 3kPa O_2_ / no serum and 1 = 37°C / 20kPa O_2_, / 10% serum, respectively. Non-parametric data (percentage live cells, total cell number, ATP measurement, qRT-PCR, DCFH-DA measurement) was analysed by means of the Mann-Whitney U test to probe for differences between two treatment groups. Results were considered to be statistically significant with * at p-values < 0.05, ** < 0.01 and *** < 0.001. Further information on individual statistical tests is available in the respective figure legends.

## Results

### Mechanical properties of tendon fascicles are impaired by *ex vivo* culture conditions mimicking vascularization

To investigate the impact of extrinsic tissue factors on underload-induced alterations in tendon core mechanical properties we established a tendon explant model. Intact tendon fascicles were isolated from murine tails and cultured deprived from load to mimic tendon underloading after damage. We cultured the tendon fascicles with or without serum and under varying temperature and oxygen levels to study the involvement of extrinsic factors associated with vascularization under tightly controlled conditions (Figure 1B). Fascicles cultured in serum-free medium, at low oxygen partial pressure (3kPa pO_2_) and low temperature (29 °C) mimic a quiescent, minimally nourished tissue that is typical of the tendon core within a healthy tissue [57–59] (Supplementary figure S1). In contrast, a pathologically vascularised tissue was mimicked by elevated temperature, oxygen and nutrients (37°C, 20kPa pO_2_, 10% serum) that would follow upon involvement of blood vessels and synovial tissue.

High tensile strength and high elastic modulus characterize a functional tendon, allowing it to transmit muscle forces across joints of the body. To evaluate the effect of tissue culture niche on tendon integrity we measured the elastic moduli of tendon fascicles after 6, 12, 18 and 24 days (d6 – d24). Elastic moduli were calculated from force-displacement curves that were recorded on a custom-made uniaxial test device [43]. The elastic modulus of freshly isolated (uncultured) fascicles was 1300 MPa (sd: 242 MPa). Under serum-free conditions, the elastic moduli were not affected by load-deprived culture and remained stable independent of oxygen partial pressure and temperature throughout the experimental period (Figure 2A, 2B). Likewise, including serum to the low temperature (29°C) and low oxygen (3kPa pO_2_) condition did not alter emergent mechanical properties (Figure 2C). In contrast, the elastic modulus was strongly and significantly decreased when maintaining the tendon fascicles under conditions that mimic a pathologically vascularized tendon niche (37°C, 20kPa pO_2_, 10% serum) (Figure 2D). Here, the elastic modulus started to decrease after day 6 and dropped to 367 MPa (sd: 426 MPa) at day 18. At day 24, we were not able to measure the elastic modulus due to a complete loss of load bearing capacity. Notably, the decrease of mechanical properties did not occur in devitalized control tendons, indicating that intrinsic cellular processes and not proteolytic activity of serum components were responsible for the decrease in elastic modulus (Supplementary figure S2).

**Figure 2:**
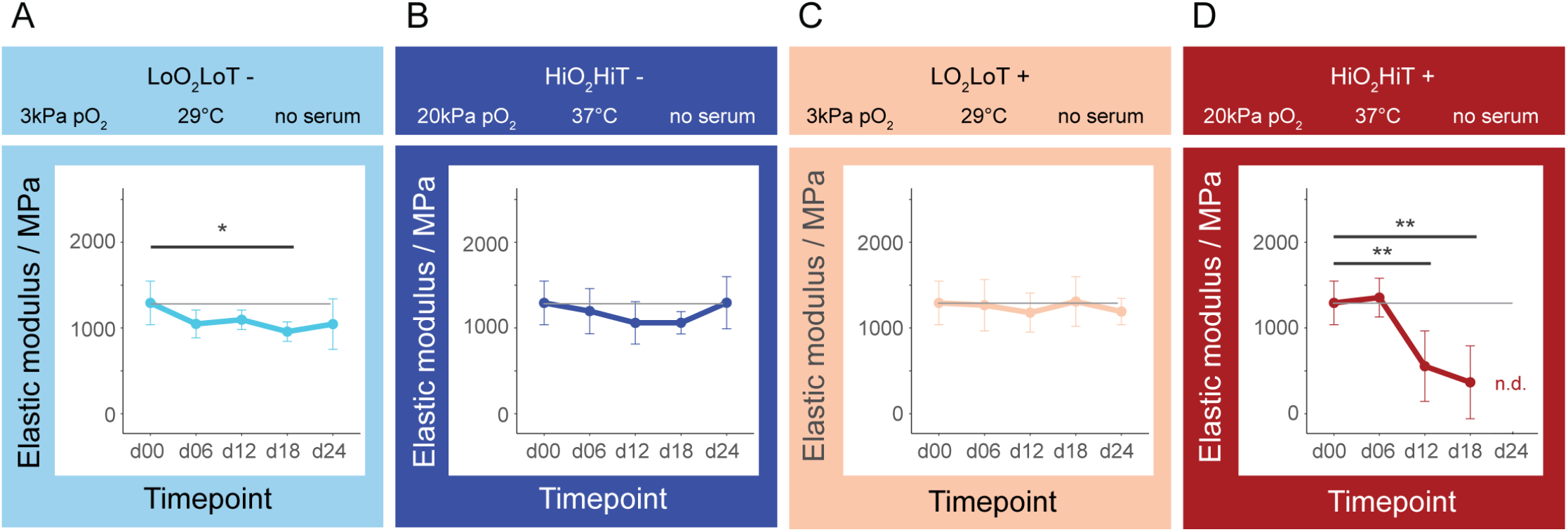
Elastic moduli of tendon fascicles cultured in different environments during 24 days (d24). The models represent increased involvement of the extrinsic compartment associated with vascularization of the tendon (**A** to **D**). Loss of mechanical properties only occurred under standard cell culture conditions in serum-supplemented medium, at a temperature of 37°C and atmospheric oxygen level (**D**). The data points represent mean values and the error bars are standard deviation. Statistical analysis was performed using one-way ANOVA followed by contrast analysis in RStudio. Mean values of each time-point (d06 – d24) were compared to the freshly isolated control (d00) within each condition (n = 6 independent samples from different mice, p-values: * < 0.05, ** < 0.01).

Oxygen- and temperature-dependent losses of mechanical properties in the serum-containing environment happened in an additive manner, whereby low oxygen (37°C, 3kPa pO_2_) and low temperature (29°C, 20kPa pO_2_) each slowed the loss of mechanical properties (Supplementary figure S3). Multiple linear regression analysis was carried out to investigate whether culture time-point, temperature, serum and oxygen predict the elastic modulus of explanted tendon fascicles. A significant regression equation was found (F(6, 127), 5.494, p = 4.37*10^−5^) with an R^2^ of 0.21. Culture time-point (β of d12 = −200.1, p = 0.02, β of d18 = −195.8, p = 6.2*10^−4^, β of d24 = −255.4, p = 6.3*10^−3^), temperature (β = −284.5, p = 2.89*10^−5^) and oxygen (β = 163.26, p = 0.013) significantly predicted the elastic modulus. Serum was not a significant predictor (β = −49.15, p < 0.45).

In summary, we established a novel *ex vivo* tendon model that represents different levels of extrinsic compartment involvement during underload-mediated tendon remodelling. The model reveals that cell-mediated processes drive loss of mechanical properties of the tendon fascicles at activating *ex vivo* conditions (high levels of oxygen and temperature).

### Comparable cell viability, metabolism and gross ECM structure between functionally impaired and intact tendon fascicles

Loss of mechanical properties in a serum-supplemented environment is driven by high temperature and oxygen in a cell-dependent manner. To gain deeper insights into the cellular processes that are responsible for loss of mechanical properties we followed up on the intact and functionally impaired groups. We therefore cultured the tendon fascicles at either low oxygen and low temperature (LoO_2_LoT+) or high oxygen and high temperature (HiO_2_HiT+) in the presence of serum for the following investigations.

To evaluate whether cell number or viability contributed to altered mechanical properties we performed a fluorescence-based cell viability assay in tendon fascicles after 6, 12, 18 and 24 days of culture. Total cell number and cell viability within the tissue were not markedly different between the two conditions as well as compared to the freshly isolated control tissue (Figure 3A). Further, metabolic activity as measured by ATP concentration normalized to DNA content increased similarly in both culture conditions after 6 and 12 days of culture compared to the freshly isolated control measured at day 0 (Supplementary figure S4). To investigate tendon fascicle morphology, we looked at macroscopic alterations and additionally performed second harmonic generation imaging of collagen. In both conditions, fascicles showed length contraction over time. Although fascicles in the HiO_2_HiT+ environment contracted at earlier timepoints compared to the LoO_2_LoT+ group, collagen fibres appeared microscopically similar in both conditions (Figure 3B). Furthermore, also the ultra-structure of collagen fibrils was qualitatively comparable between both groups as revealed by transmission electron microscopy (Figure 3B).

**Fig. 3:**
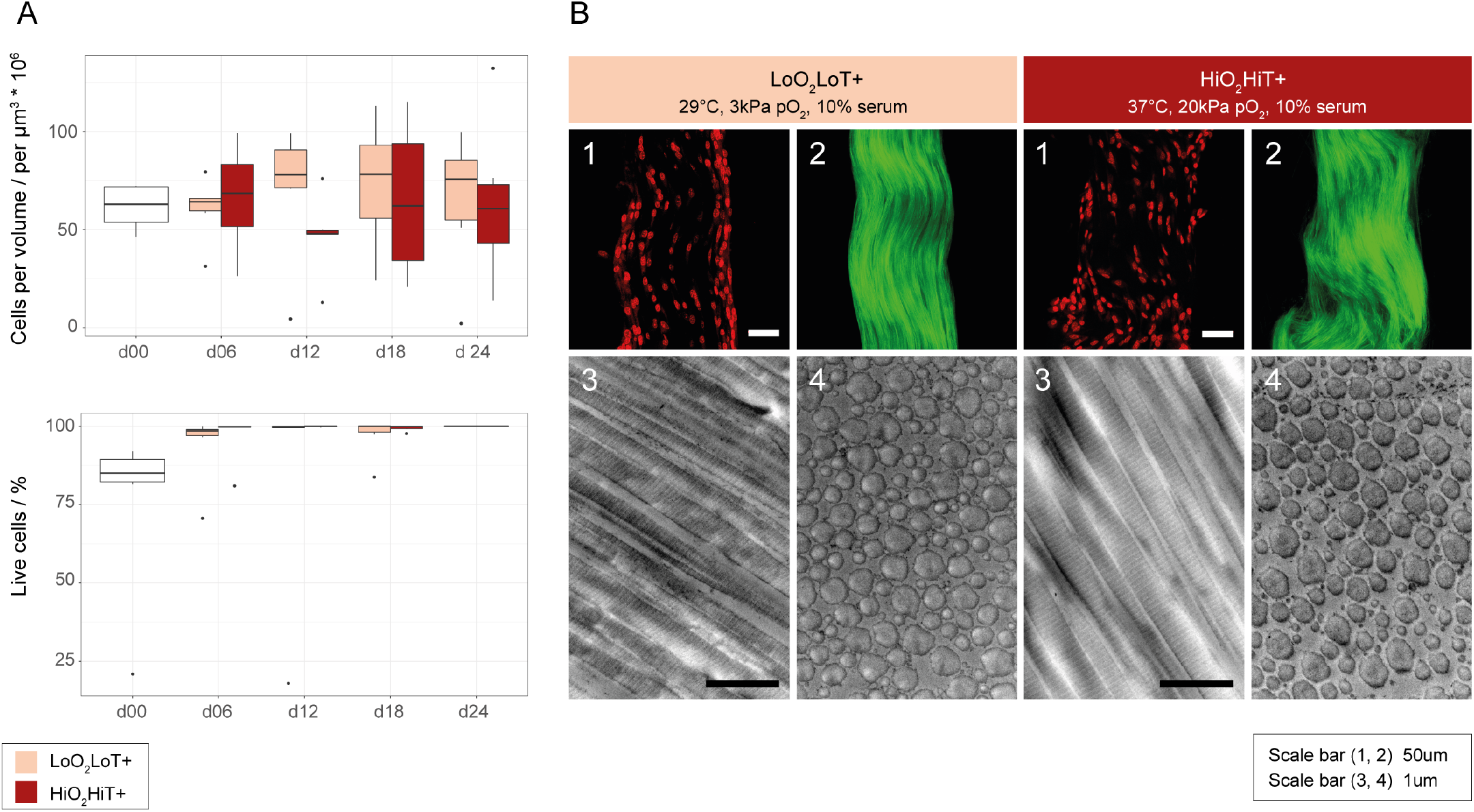
Cell viability and morphological characterization of mechanically intact and impaired tendon fascicles. (A) Total cell number per volume (upper) stayed constant over time and cell viability (lower) was not significantly different between the HiO_2_HiT+ and the LoO_2_LoT+ group at any time-point. Single dead peripheral cells were always present in the d0 condition, which most probably died from shear forces occurring during the isolation process. However, these cells disappeared upon culture in serum, which explains the slightly lower percentage of live cells in the d0 condition. Statistical analysis was performed using Mann-Whitney U test in RStudio comparing the “LoO_2_LoT+” to the “HiO_2_HiT+” condition within each time point (d06 – d24) (n = 6, independent samples from different mice). (B) Tissue morphology after a treatment period of 12 days. (1) Second harmonic generation imaging and (2) nuclear shape of mouse tail tendon fascicles treated at either low oxygen and low temperature with serum (LoO_2_LoT+) or high oxygen and high temperature with serum (HiO_2_HiT+). (3) Transmission Electron Microscopy images in longitudinal and (4) transverse sections. Scale bars represent 50μm in the upper fluorescence images and 1um in the electron micrographs.

Taken together, our data show that the loss of mechanical properties does not correlate with pronounced changes at the cell or tissue matrix level. Therefore, we next aimed at investigating changes that take place at the molecular level.

### Mechanical impairment is associated with strong transcriptome changes related to immune system response, oxidative-stress, lysosome activation and proteolysis

To investigate the molecular changes that are involved in the loss of mechanical properties we compared the functionally impaired (HiO_2_HiT+) and intact (LoO_2_LoT+) tendon fascicles by transcriptome analysis using RNA sequencing (RNA-seq). Overall we detected 23’675 known transcripts according to the murine reference genome. Filtering revealed 3305 differentially expressed genes (DEG) between the two groups, among which 1854 were up-regulated and 1451 were down-regulated in the functionally impaired fascicles (Figure 4A and supplementary table ST1). To systematically investigate their relationships, we functionally enriched all the DEGs within the gene ontology (GO) Biological Process. Overrepresented GO terms mainly include immune-related processes (Supplementary table ST2), whereof nearly 50% are additionally represented in the GO enrichment of only up-regulated genes (Supplementary table ST3). Furthermore, we mapped all DEGs onto the most relevant pathways via statistical enrichment analysis using the MetaCore database by GeneGo. The 10 most significantly altered pathways are shown in Figure 4B. Strikingly’ 7 out of 10 pathways are related to immune system activation including professional phagocytic cells’ such as neutrophils’ macrophages and dendritic cells. In addition’ three identified pathways (“Chemokines in inflammation”‘ “HIF-1 target transcription” and “Activation of NADPH oxidase”) are closely linked to oxidative-stress-dependent processes (Figure 4B). The induced oxidative-stress related genes mainly include regulatory proteins or subunits of the NADPH oxidase (NOX2/gp91^phox^’ p22^phox^’ p47^phox^’ p67^phox^’ p40^phox^’ Rac2’ Vav1’ Vav3)’ a transmembrane protein that catalyzes the production of superoxide in different cell types (Supplementary figure S5A). All of these genes were significantly up-regulated in the HiO_2_HiT+ group.

**Figure 4:**
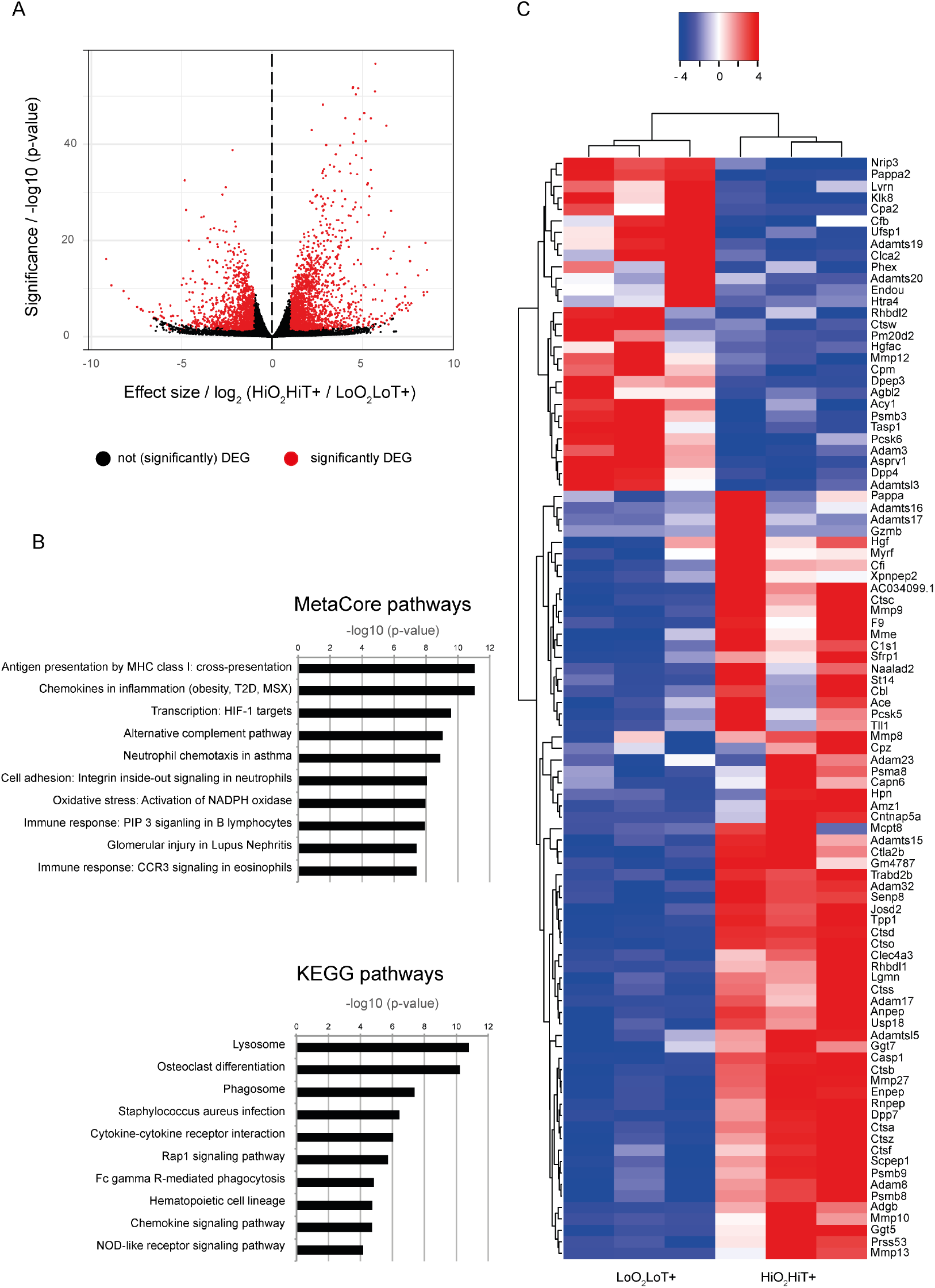
Transcriptome analysis of differentially expressed genes in mechanically intact vs. impaired tendon fascicles. (A) Volcano plot of differentially expressed genes between tendon fascicles cultured in HiO_2_HiT+ and LoO_2_LoT+. Genes coloured in red are considered as differentially expressed (DEG) and have a log_2_(fold change) > 1 and a significance of p < 0.05. (B) The top 10 enriched signalling pathways in the HiO_2_HiT+ group compared to the LoO_2_LoT+ group (determined by the MetaCore from GeneGo and KEGG database using both, up- and down-regulated genes). MHC = Major Histocompatibility Complex, T2D = Type 2 Diabetes, MSX = Metabolic Syndrome X (C) Unsupervised hierarchical clustering of differentially expressed genes in the LoO_2_LoT+ and the HiO_2_HiT+ group (n=3, independent pools of fascicles from different mice). Depicted are the DEGs with cut-off values of log_2_ ratio > 1 and p < 0.05. The clustering separates the genes by colour with positive or negative row-scaled Z-scores represented in red and blue, respectively.

Alternatively, we enriched all DEGs in the Kyoto Encyclopedia of Genes and Genomes (KEGG) domain. One half of the top 10 identified gene pathways is associated with inward vesicular trafficking destined to degradation in the lysosome (“Lysosomes”, “Phagosomes”, “Fc gamma R-mediated phagocytosis”, “Staphylococcus aureus infection”, “NOD-like receptor signaling pathway”) (Figure 4B). The lysosome is the main catabolic subcellular organelle responsible for degradation of extracellular and intracellular components. Furthermore, “Osteoclast differentiation” suggests initiation of a process that is relevant in ECM resorption as shown by up-regulation of the genes PPARG, CSF-1, PI3K, RANK (TNFRSF11A) and RANKL (TNFSF11) and others (Supplementary figure S5B). The KEGG pathway analysis including only up-regulated DEGs is mostly consistent with the analysis including both, up- and down-regulated genes (Supplementary figure S5C).

Because we previously showed that tendon fascicles in the HiO_2_HiT+ condition lose mechanical properties, which most likely involves ECM breakdown, and due to the fact that pathway analysis of DEGs suggests lysosomal degradation as a main process, we examined the GO term “proteolysis” (GO:0006508). Indeed, we found 96 DEGs to be annotated to “proteolysis” of which 67 were up-regulated in the HiO_2_HiT+ (Figure 4C). Among them we found various extracellular proteases such as matrix metalloproteinases (MMP 8, 9, 10, 13, 27), intracellular proteases involved in lysosomal degradation such as cathepsins (cathepsin A, B, C, D, F, O, S, Z), and proteasome-related proteins (PSMA 8, PSMB 3, 8, 9,).

In summary, the results reveal that the transcriptome changes between the mechanically intact (LoO_2_LoT+) and the mechanically impaired (HiO_2_HiT+) group involve proteolysis, oxidative-stress-related processes, immune system activation and lysosome involvement. These molecular changes are all accompanied by a functional loss of fascicle mechanics.

### Tendon mechanical integrity is rescued by inhibiting reactive oxygen species and proteases

Based on the experimental design (difference in temperature and oxygen saturation between the two groups) and the up-regulated oxidative stress-related pathways we hypothesized that reactive oxygen species (ROS) contribute to ECM degradation and loss of tendon mechanical properties as main regulators. To confirm ROS production we measured fluorescence of dichlorofluorescein (DCF), which is produced by intracellular oxidation upon esterase cleavage of the cell-permeable 2’-7’-Dichlorodihydrofluorescein diacetate (DCFH-DA). ROS were significantly increased in the functionally impaired (HiO_2_HiT+) compared to the intact (LoO_2_LoT+) tendon fascicles, confirming the RNA-seq findings on the tissue level (Supplementary figure S6).

Next, based on our RNA-seq data we hypothesised that ROS and MMPs are involved in the process of tendon fascicle degeneration. Therefore, we assessed the effects of ROS scavenging (Tempol) and MMP inhibition (Ilomastat) in the functionally impaired (HiO_2_HiT+) condition over a time period of 12 days. Separate inhibition of ROS and MMPs partially protected against loss of mechanical properties as measured by the elastic moduli (Figure 5A). Therefore, we also investigated the combined effects of inhibiting both ROS and MMPs. Intriguingly, combined inhibition of ROS and MMPs resulted in complete maintenance of elastic modulus at levels of freshly isolated control fascicles. A passive chemical cross-linking effect of the inhibitory substances could be excluded by control experiments on devitalized tendon tissue samples (Supplementary figure S7). Based on the additive effect of blocking ROS and MMPs we hypothesized that ROS and MMPs may be hierarchically linked. To address this hypothesis we performed RT-PCR on fascicles cultured in the HiO_2_HiT+ environment with or without Tempol-treatment blocking ROS. We examined eight genes of interest that had been previously related to proteolysis in our RNAseq results (MMP9, 10 and 13, cathepsin B, C, D, S and Z). Gene expression of MMP9, 10 and cathepsin C was significantly down-regulated by inhibiting ROS with Tempol after 12 days (Figure 5B). A trend towards lower expression of MMP13, cathepsin B and S was observed too. Expression of cathepsins D and Z was not affected by inhibition of ROS. These data show that ROS regulates MMP and cathepsin gene expression directly or indirectly through factors up-stream of nuclear transcription of MMPs.

**Figure 5:**
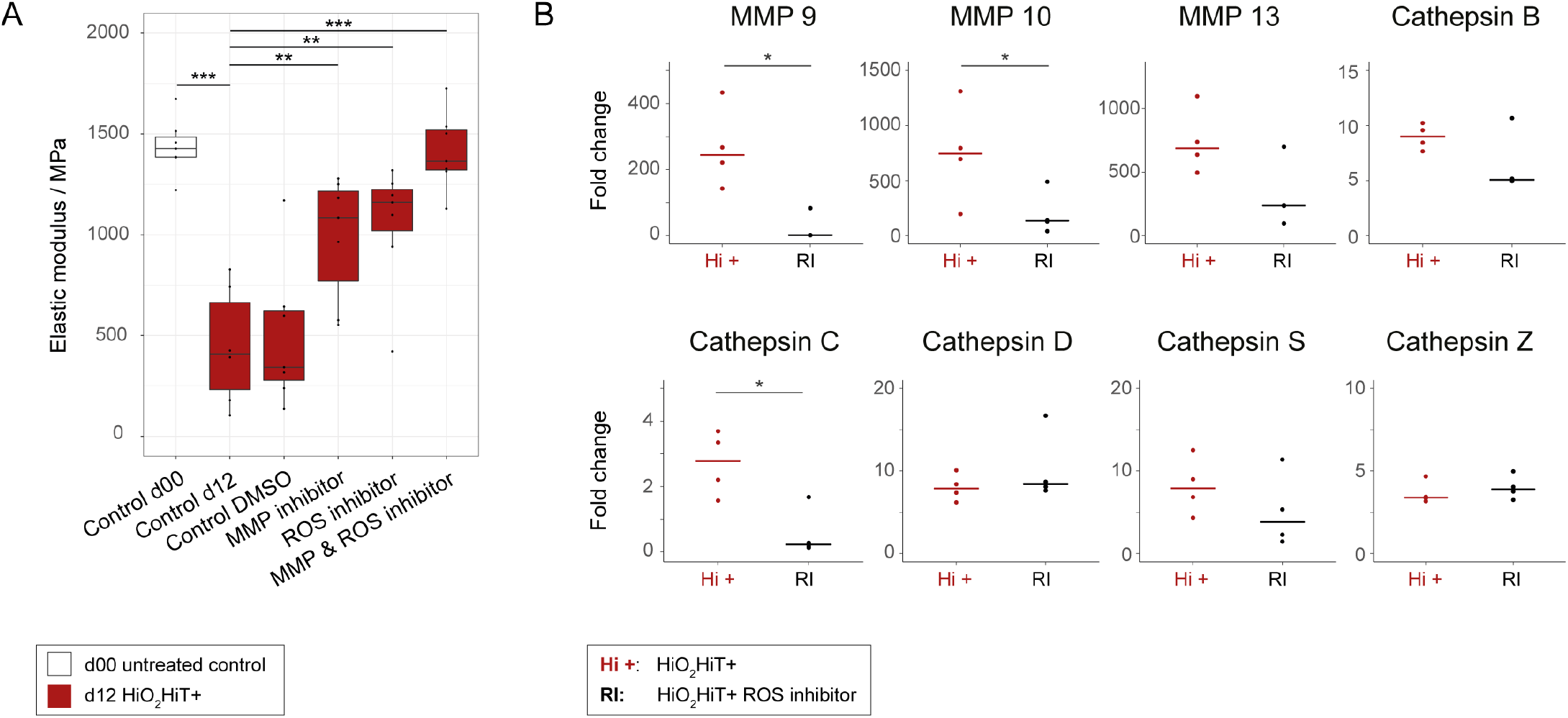
Inhibition of reactive oxygen species (ROS) and MMPs in mechanically impaired tendons. (A) Inhibition of ROS and matrix metalloproteinases by Tempol and Ilomastat, respectively. After a culture time of 12 days inhibition of ROS or MMP result in increased elastic modulus compared to the control group. Inhibition of ROS and MMPs show an additive effect restoring original mechanical properties to levels of freshly isolated tissue (Control d0). Statistical analysis was performed using one-way ANOVA followed by contrast analysis in RStudio. Mean values of groups treated with inhibitory substances were compared to the control group cultured for 12 days in serum-supplemented medium (Control d12) (n = 6). (B) RT-PCR gene expression of proteins involved in extracellular matrix remodeling after 12 days of culture with and without the ROS inhibitor Tempol. MMP 9, MMP 10 and cathepsin C were significantly down-regulated (p-value = 0.029 for all) in tendon tissue by inhibiting ROS. Gene expression of MMP 13 and cathepsins B, D, S and Z was not significantly affected by ROS inhibition. However a trend of regulation towards native gene expression is observed in MMP 13 and cathepsin B and S. Displayed are the relative fold changes compared to the freshly isolated control at day 0 and the horizontal crossbar represents the median. All values were normalized to the reference gene (Anxa5). Mann-Whitney U tests were performed using in RStudio by comparing the Tempol-supplemented group to the “HiO_2_HiT+” group (n = 4, p-values: * < 0.05, ** < 0.01, *** < 0.001).

Taken together, our findings suggest that elevated oxygen and temperature act as pathological drivers of fibroblasts in the tendon core. These cells undergo multifactorial molecular processes that involve oxidative-stress and immune system activation pathways. Inhibiting these pathways was sufficient to abrogate extracellular matrix degradation and functional impairment in unloaded tendon explants.

### Tendon degradome shows altered activity of matrix metalloproteinases and lysosomal proteases and increased collagen type I cleavage

To quantify proteins and N termini and thereby identify proteolytic cleavage events in intact and degraded tendon explants, we performed 10plex-Tandem Mass Tag^TM^-Terminal amine isotopic labeling of substrates (TMT-TAILS) (Figure 6A). 382 proteins were quantified with high confidence (FDR < 0.01), of which 35 were significantly more and 13 significantly less (adj. p-value < 0.05; fold change > 1.5) abundant in the degraded HiO_2_HiT+ phenotype group (Figure 6B, Supplementary table ST4). More abundant hits included proteins associated with proteolysis or lysosomes (MMP3, MMP13, beta-glucuronidase, lysosomal protective protein/cathepsin A, cathepsin Z, V-type proton ATPase subunits A/E1/G1, lysosomal acid phosphatase), intracellular trafficking (sorting nexin 3/5, programmed cell death 6-interacting protein, glycolipid transfer protein) and ROS metabolism (catalase, peroxiredoxin 1). Less abundant proteins are associated with cytoskeletal organization (vimentin, stathmin, cochlin, myocilin, plectin, moesin) or cell-cell and cell-matrix interaction (thrombospondin 1/4).

**Figure 6:**
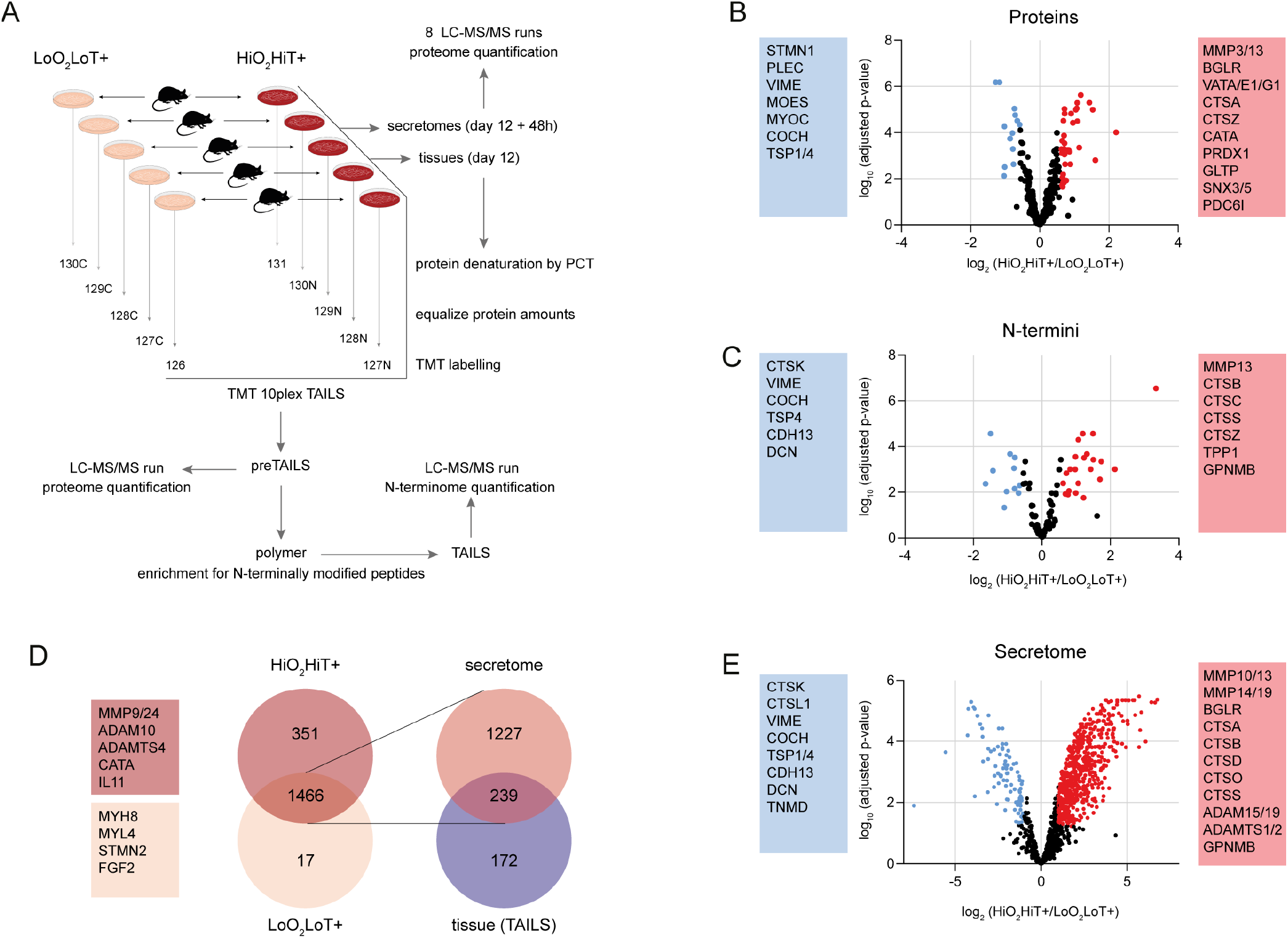
Proteomic analysis mechanically intact and impaired tendon fascicles. (**A**) TMT-TAILS workflow. Tissues of each condition (n=5, independent pools of fascicles from different mice) were collected after 12 days. Protein amounts of all tissue samples were equalized and subsequently analyzed in a 10plex TMT-TAILS experiment. Medium for secretome (n=4) collection was additionally conditioned for 48 hours and analyzed by mass spectrometry based label-free protein quantification method. (**B**) Quantified proteins and (**C**) N termini from TMT-TAILS probes. Proteins coloured in red and blue are considered as differentially more or less abundant and have significance of p < 0.05 and a log_2_(fold change) > 1.5 (red) or < 1.5 (blue). (**D**) Venn diagram of quantifiable proteins in the secretomes of the mechanically intact (LoO_2_LoT+) and impaired group (HiO_2_HiT+). 239 out of totally 1466 secreted proteins were detected both extracellular (released into the medium) and intracellular (tissue, TAILS experiment). (**E**) Label-free quantification of the 1466 secreted proteins of which 860 show significantly different abundance. Red dots (740) show significantly more (p < 0.05, log_2_(fold change) > 2) and blue dots (120) significantly less (p < 0.05, log_2_(fold change) < 2) abundant proteins in the HiO_2_HiT+ group. ADAM = a disintegrin and metalloproteinase, ADAMTS = a disintegrin and metalloproteinase with thrombospondin motif, BGLR = beta-glucuronidase, CATA = catalase, CDH = cadherin, COCH = cochlin, CTS = cathepsin, MMP = matrix metalloproteinase, DCN = decorin, FGF = fibroblast growth factor, GLTP = glycolipid transfer protein, GPNMB = transmembrane glycoprotein NMB, IL11 = Interleukin 11, MOES = moesin, MYH = myosin, MYL = myosin light chain, MYOC = myocilin, PDC6I = programmed cell death 6-interacting protein, PLEC = plectin, PRDX = peroxiredoxin, SNX = sorting nexin, STMN = stathmin, TNMD = tenomodulin, TPP1 = lysosomal pepstatin-insensitive protease, TSP = thrombospondin, VAT = V-type proton ATPase subunits, VIME = vimentin.

Additionally, we extracted 103 N-terminally TMT-labeled N-terminal semi-tryptic quantifiable peptides from 79 different proteins, whereof 46 could be assigned to curated protein start sites and 57 were protease-generated neo-N termini (Figure 6C, Supplementary table ST5). Quantitative N-terminome analysis revealed significantly (adj. p-value < 0.05; fold change > 1.5) increased abundance of active lysosomal proteases (cathepsins B, C, S, Z, lysosomal pepstatin-insensitive protease) all identified by their mature protein N terminus generated upon propeptide removal in the degraded tendon tissue, whereas to our surprise the mature protein N terminus of cathepsin K was inversely abundant between conditions. Transmembrane glycoprotein NMB, which has been identified in osteoclast and macrophage differentiation, was highly abundant in the degraded phenotype and could be determined by a cleavage site within its extracellular domain. Moreover, N-terminomics corroborated increased abundance of MMP13 in the same group, but by identification of a neo-N-terminal peptide, indicating removal of the hemopexin domain with potential implication in altered substrate specificity. Although below the fold-change cut-off of 1.5, we identified 10 collagen type I cleavage sites, of which 8 were significantly more abundant in the mechanically degrading group. Four of these cleavage products were associated with the collagen type I alpha 1 chain (Col1a1) including a known MMP13 site and 4 with the alpha 2 chain (Col1a2).

We then quantitatively compared proteins released into supernatants of *ex vivo* tendon cultures under degrading and non-degrading conditions. We identified a total number of 2017 proteins with high confidence (FDR < 0.01) (Supplementary table ST6), of which 351 were only found in the supernatants of the HiO_2_HiT+ group (quantifiable in at least two replicates) (e.g. catalase, many proteasome proteins, MMP9 and 24, ADAM10, IL11) and 17 only in the LoO_2_LoT+ group. Differential abundance was calculated from 1466 proteins, of which 226 proteins were identified also in the tissue sample (Figure 6D). 740 proteins were significantly more abundant (adj. p-value < 0.05; fold change > 2) in the secretome of the degraded group, accounting for ~85% of all 860 proteins with significantly differential abundance. Quantification of the released proteins corroborated the protease abundance pattern of the tissue proteome showing significantly increased extracellular secretion of proteases, such as cathepsins A, B, D, O, S and MMP13, 14 19, in the degraded group (Figure 6E), while cathepsin K was significantly less abundant in the supernatants of the HiO_2_HiT+ group.

## Discussion

Tendon is a tissue with a low innate regenerative capacity, a limited number of resident fibroblasts with normally low metabolic activity, and low tissue vascularity [60, 61]. The mechanisms of tendon remodelling after injury or in chronic tissue pathologies are poorly understood. Observational studies describe the recruitment of cells from the extrinsic tissue compartments (blood vessels, inflammatory mediators) to the tendon core in chronically disordered tendon tissue [39–42]. However, the interplay between the intrinsic tendon core comprised of collagenous ECM and tendon fibroblasts, and the extrinsic peritendinous support tissues during injury and repair remains largely unknown. In this work we studied how the tendon core responds to mechanical load deprivation (that mimics a damage scenario), and how this response may be mediated by presence or lack of a vascularized tissue niche. This question has implications to both understanding chronic tendon disease, and surgical repair of torn tendons.

In this study, tendon microtear with localized matrix unloading was modelled using free-floating culture of murine tail tendon fascicle explants. The fascicle *ex vivo* explant model maintains the cells in their native tissue microenvironment, allows normal cell-cell and cell-matrix communication, and is well suited to the mechanistic study of cell-mediated matrix remodelling in pathophysiological contexts [62–66]. Exploiting this model, we confirmed that standard tissue cultures (elevated nutrient supply, 37°C and 20kPa pO_2_) triggered the loss of mechanical properties as shown previously [63, 64, 67, 68]. Strikingly, tendon fascicles that were cultured free-floating with serum at 29°C and 3kPa pO_2_ fully maintained their mechanical properties as well as their viability. The additional extrinsic cues of hyperoxia and hyperthermia were required to initiate stromal compartment remodelling. We interpret these cues to represent a vascularized tendon niche, such as expected after severe injury or in advanced stages of tendon disease.

After verifying that passive thermodynamic effects of elevated tissue culture on proteolytic kinetics were not responsible for accelerated tissue degradation at higher temperatures [69] (Supplementary figure S3), we investigated the underlying cellular mechanisms. With no obvious differences in macro-nor microscopic tissue structure in conditions to explain the functional loss of tissue strength, we performed a deeper molecular investigation. Comparative quantitative degradomic analysis revealed measurable cleavage of collagen type I in the degraded tissue, consistent with the according loss of macro-mechanical properties. Furthermore, proteomics data showed a general increase in protein release to the supernatant in the functionally impaired group, indicating an elevation in extracellular signalling and secretory capacity under pathological conditions. Activation of cathepsins and MMPs in the degraded phenotype strongly supported the transcriptomic analysis, elaborating in detail the widespread elevation of protease expression. By RNAseq we found that ROS and induction of matrix proteases (MMPs and cathepsins) are linked, which has been widely described earlier in vascular, cardiac, bone and cancer tissues [70–73]. Although ROS have been occasionally mentioned in tendon-related literature their link to protease activation in tendon remodelling processes has been largely overlooked [74–76]. While we can conclude from our data that ROS lay upstream of protease activation, it remains unknown whether hyperthermia and hyperoxia activate a precursor component in the serum that serves as an ignition switch for parallel or consecutive processes, such as ROS generation, tissue degradation and immune system activation. Early research on muscle suggests that hyperthermia induced by exercise directly causes oxidative stress through ROS generation, changes in the immune system and inflammatory response by increased cytokine production [77–79]. According to our RNA-seq dataset it is also conceivable that ROS production and subsequent MMP activation occur via NADPH oxidase. We identified the CYBB gene (also known as NOX2), a member of the NADPH oxidase family, which is described to be present in a large number of tissues including thymus, colon, testis, neurons, skeletal muscle myocytes and endothelial cells, among others [80]. However, it is still widely accepted that CYBB has phagocyte-specific tissue expression. Therefore it seems likely that the cells of the tendon fascicles express a phagocytic-like character. This assumption is further strengthened by the identification of different pathways that are associated with phagocytosis and lysosomes. Phagocytosis is a process of endocytosis when extracellular solid particles are engulfed. The phagosome, the intracellular vacuolar compartment generated during phagocytosis, fuses with the lysosome, an intracellular structure that contains hydrolytic enzymes such as cathepsins, to produce the phagolysosome [81].

Furthermore, we identified induced pathways and proteins widely associated with differentiation of osteoclasts, cells closely related to the family of professional phagocytes [82]. Specifically, the up-regulated PPAR gene, which is involved in osteoclast differentiation processes, was recently shown to induce a lysosome-controlled ECM catabolism [83]. In line with these results, matrix resorption in our degraded tendon model might be accelerated by lysosomal proteases upon release from lysosomal vesicles that fuse with the plasma membrane as cathepsins were found both in tissue proteome and secretome [84]. While in bone, cathepsin K is the driving force in ECM degradation, our degradomic data suggest that cathepsins B, C, S and Z are mainly involved in tendon tissue breakdown [85]. Interestingly, many cells with a phagocytic character are also the effector cells of the innate immune system thereby acting to enhance inflammation. Likewise, fragments of proteases may also elicit pro-inflammatory signalling [86–88]. Therefore, it is tempting to speculate that phagocytosed ECM components are cleaved by proteases within the lysosome and the generated fragments may subsequently serve as mediators of inflammation and immune response as so called damage-associated molecular patterns (DAMPs) [89–91]. DAMPs are known to initiate a non-infectious inflammatory response by activating antigen-presenting cells to become stimulatory to the adaptive immune system [92, 93].

Taken together, exogenous peptides generated from extracellular matrix breakdown after acute loss of mechanical tension in a pathological tissue may be internalized by phagocytosis, processed in the phagolysosome to produce fragments that serve as “antigens”, which are finally presented to the immune system via class I MHC (also found induced in our RNA-seq data) in tendon fibroblasts. This process, also known as cross-presentation [94], has been identified as the most significantly regulated pathway in our impaired tendon model. It may be that a small fraction of cleaved matrix fragments remain to act as DAMPs, while the majority is eventually recycled into the functional matrix. This concept of recycling of ECM proteins during remodelling is supported by evidence that *de novo* synthesis of ECM in healthy tendon appears to be very limited over the course of an individual lifetime [95].

Among the limitations of the present work is that it only captures unidirectional extrinsic gating of the intrinsic compartment, neglecting bidirectional signaling between the intrinsic and extrinsic tissue compartments. Interpreting how signals released into the extracellular microenvironment act on cells and ECM might be instructive for further deciphering the communication between tissue compartments in the context of immune response, for better understanding the pathological mechanisms and revealing a possible phenotype shift of cells within the native tissue.

Overall our data demonstrate that tissue temperature and oxygen levels represent decisive contextual cues in triggering ECM proteolysis after mechanical unloading. Hyperoxic and hyperthermic conditions mimic pathological vascularity, and apparently act as a gate to the onset of catabolic tissue turnover. The involved pathways include ROS, protease induction and immune-cell recruitment. Our data provide key insights into how initially quiescent tendon cells are activated to work as executers of tissue remodelling in the absence of active extrinsic helper cells. A major aspect of our work is that it highlights the physiological role of low oxygen and below body core temperatures as important in maintaining cellular quiescence and functional integrity of the ECM. Mechanical tension is dogmatically considered to be essential to protect tendon tissue from proteolytic turnover and tissue remodelling. It is therefore remarkable that mechanics alone were insufficient to trigger proteolytic matrix turnover in low oxygen and temperature. Among other implications, our findings suggest that tendon cell and tissue culture conditions should be reconsidered in experimental settings that study healthy tissue. This does not only apply for the tendon field but also for scientific investigations of other stromal tissues with normally low vascularity, such as cartilage, the intervertebral discs and ligaments.

We present a tendon *ex vivo* model that by changing the culture environment can be used to reflect tendon physiology or pathophysiology. To our knowledge this is the first study to in-depth investigate the events leading to tissue degradation at the tissue (biomechanics, tissue morphology, cell viability) and molecular level (transcriptomics and degradomics) and thereby offering broad insight into how multifactorial molecular processes may converge to a possible mechanism, by which ECM homeostasis is shifted towards degradation. Collectively, we anticipate that our study will have a significant impact in the field of tendon biology by contributing to a more mechanistic understanding of tendon physiology and at the same time helping to effectively progress the development of effective treatments for tendinopathy. Furthermore, our observations give rise to many new hypotheses for future investigations involving ECM remodelling, oxidative stress and immune-related cellular response in tendon and stromal tissue.

## Acknowledgment

We gratefully thank Max Hess and Lucien Segessemann for their important contribution and expertise in image analysis, Jonathan Ward, Susanne Freedrich and Kerstin Sacher (ETH Phenomics Center) for providing the research animals and their assistance in the animal facility. We thank Ursula Lüthi and Andrés Käch (Center for Microscopy and Image Analysis, Zürich) for TEM microscopy, Lilian Hartmann and Olivier Leupin for technical assistance in the RNA isolation procedure and Jelena Kühn Georgijevic and Weihong Li (Functional Genomics Center, Zürich) for their support on RNA sequencing data and Tobias Götschi for the support on statistical data analysis. This work has been funded by ETH (Grant 12-13-2), the Vontobel Foundation, the Swiss National Science Foundation (P1EZP3-181729) and a Novo Nordisk Foundation Young Investigator Award (NNF16OC0020670).

## Author contributions

Study concept and design: SLW, FP, US, JGS. Collection and/or assembly of data: SLW, UB, ABP, BN, LB. Data analysis: SLW, UB, CNH, UADK. Data interpretation and manuscript writing: SLW, UB, UADK, JGS.

## Supplementary figures

**Figure S1:**
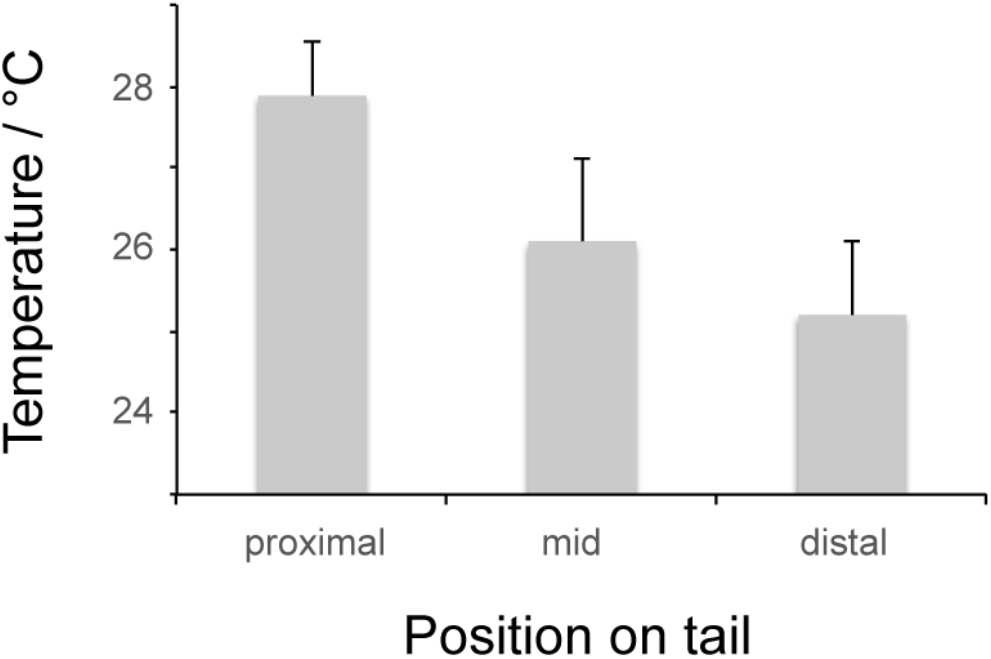
Superficial temperature of the mouse tail was estimated by measuring infrared emission at different locations. Skin temperature decreases with increasing distance to the body core. The tail of rodents is thought to act as a temperature regulator. Heat is brought by the blood flow through the central tail artery and is dissipated through the skin. The columns represent mean values and the error bars standard deviation (n = 40).

**Figure S2:**
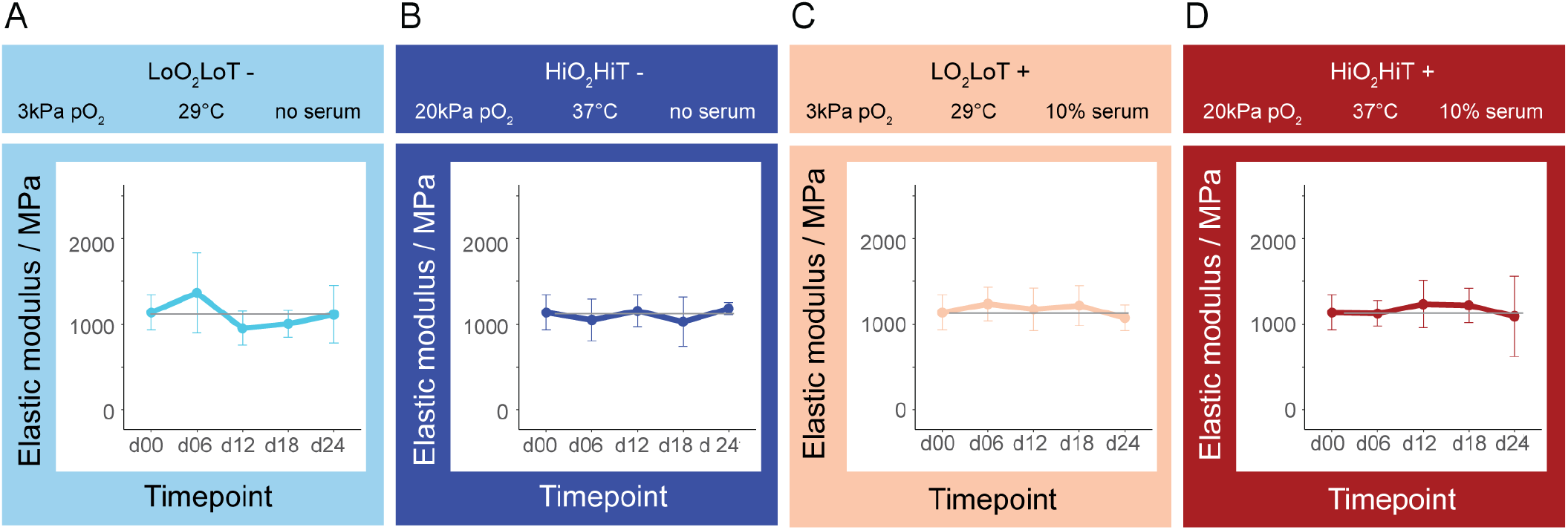
Elastic moduli of devitalized tendon fascicles cultured in the different environments over a time period of 24 days (d24). The data points represent mean values and the error bars represent standard deviation. Statistical analysis was performed using one-way ANOVA with a contrast matrix in RStudio. Mean values of each time-point (d06 – d24) were compared to the freshly isolated control (d00) within each condition (n = 6 independent samples from different mice).

**Figure S3:**
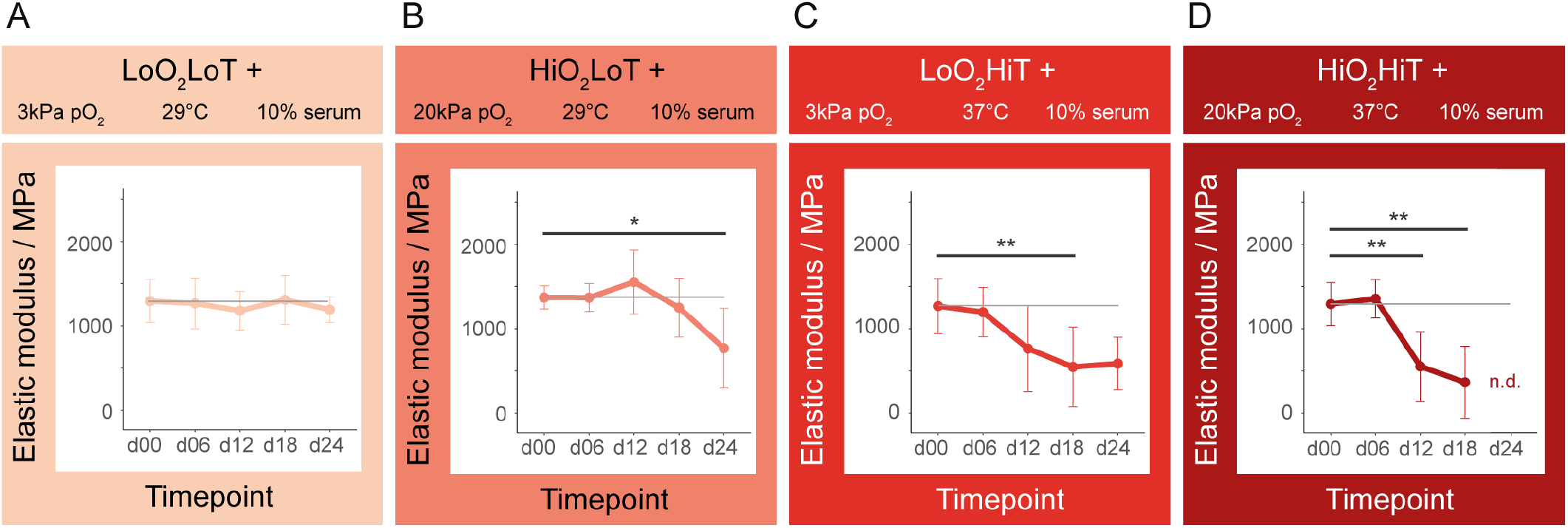
Elastic moduli of tendon fascicles cultured in different environments during 24 days (d24). The models represent extrinsic compartment involvement with progressive increase in severity (**A** to **D**). The additional culture conditions were introduced to better understand the quasi-decoupled effect of high temperature or high oxygen on extracellular matrix degradation. The data points represent mean values and the error bars are standard deviation. Statistical analysis was performed using one-way ANOVA followed by contrast analysis in RStudio. Mean values of each time-point (d06 – d24) were compared to the freshly isolated control (d00) within each condition (n = 6 independent samples from different mice, p-values: * < 0.05, ** < 0.01).

**Figure S4:**
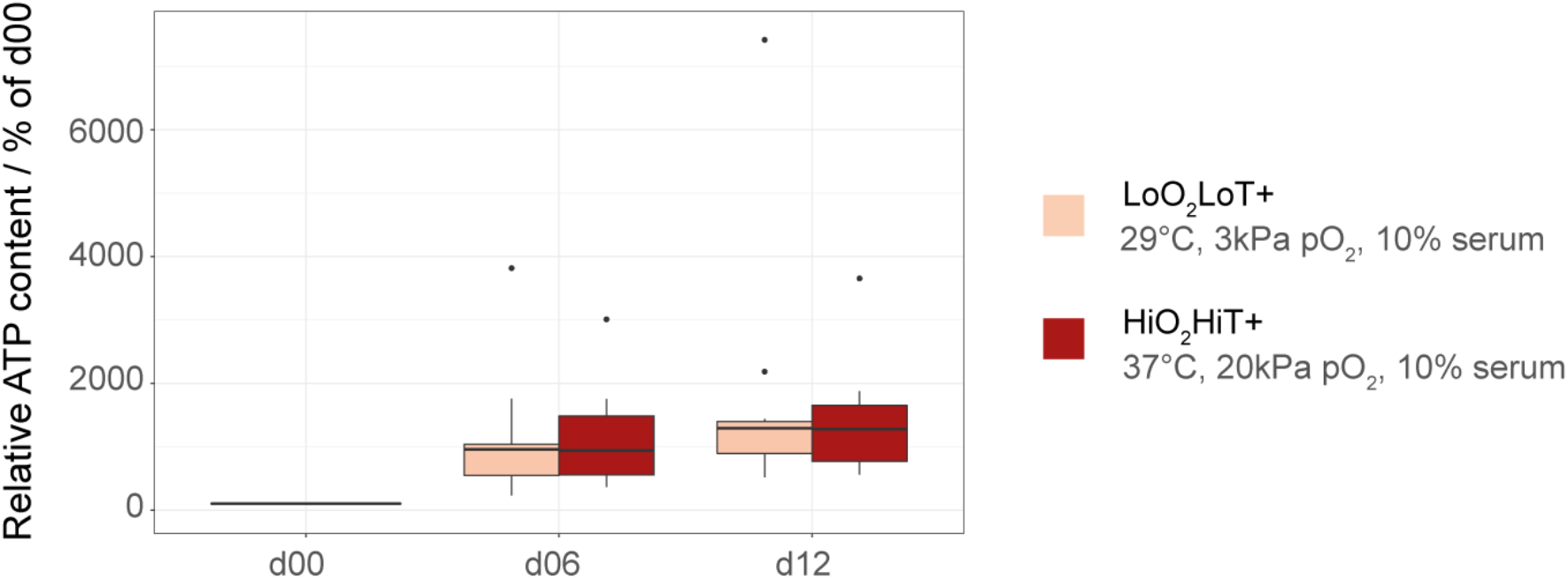
Normalized ATP content of mechanically intact and degraded tendon fascicles. Normalized ATP content was comparable between groups at each timepoint (d06 and d12). Boxplots display median value and 1^st^/3^rd^ quartiles. Outliers are plotted as individual data points. Three outliers > 7500 were removed. Statistical analysis was performed using Mann-Whitney U test in RStudio comparing the “LoO_2_LoT+” to the “HiO_2_HiT+” condition within each time point. (n = 4 in duplicates, with two measurements per animal).

**Figure S5:**
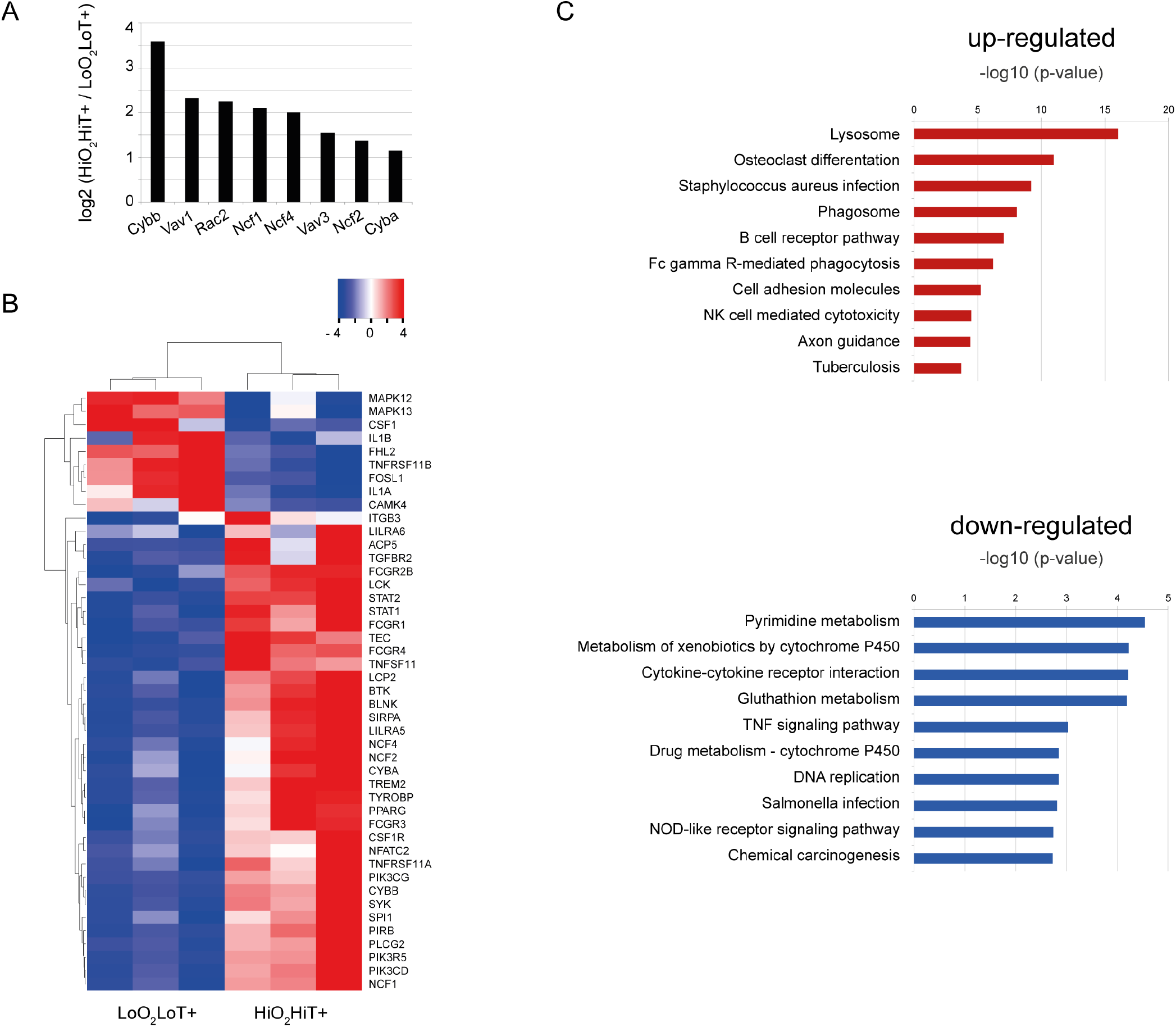
RNA-seq data analysis of differentially expressed genes comparing mechanically intact and degraded tendon fascicles. (**A**) All genes associated with NADPH oxidase are significantly up-regulated in the functionally impaired group (HiO_2_HiT+). Represented are the significantly differentially expressed genes with a log2 ratio of > 1 and a p-value < 0.05. (**B**) Heatmap of genes mapped to the KEGG pathway “Osteoclast differentiation”. The cluster separates the differentially expressed genes by colour with positive or negative row-scaled Z-scores represented in red and blue, respectively. (**C**) The top 10 enriched signalling pathways in the HiO_2_HiT+ group compared to the LoO_2_LoT+ group determined by the KEGG database separated by up- and down-regulated genes.

**Figure S6:**
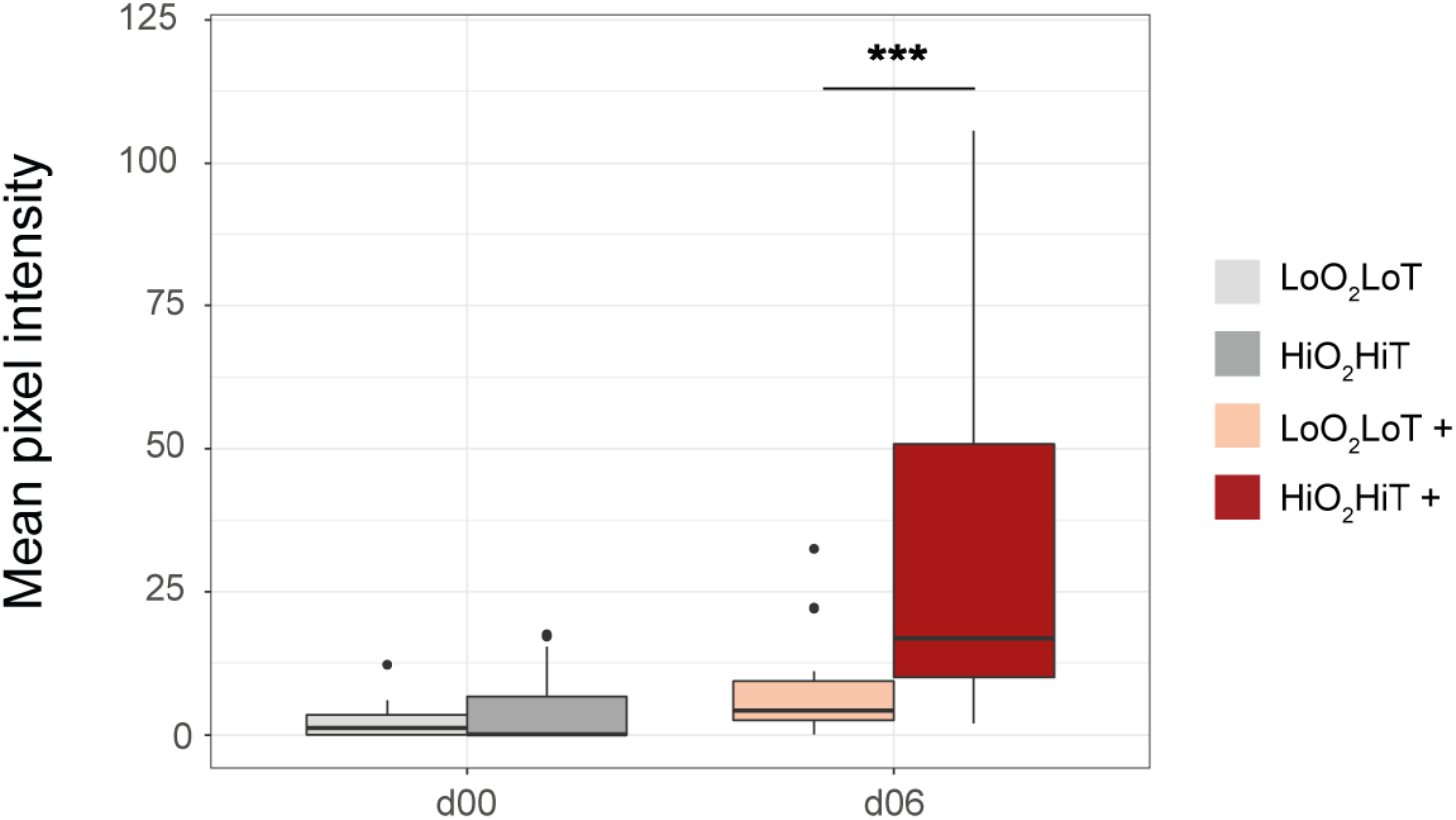
Detection of reactive oxygen species (ROS) by quantification of fluorescence intensity of DCF in tendon fascicles upon incubation at either 29°C and 3kPa pO_2_ (LoO_2_LoT) or 37°C and 203kPa pO_2_ (HiO_2_HiT) for 30min directly after extraction from the tail (d00) or after incubation in serum containing medium (+) in LoO_2_LoT or HiO_2_HiT environment for 6 days. Boxplots display median value and 1^st^/3^rd^ quartiles. Mann-Whitney U test (RStudio) was used to probe statistically significant differences between the LoO_2_LoT and HiO_2_HiT condition within each time point. (n = 6, *** < 0.001). Presence of ROS was significantly different between the two culture environments at day 6 (p-value = 0.0004).

**Figure S7:**
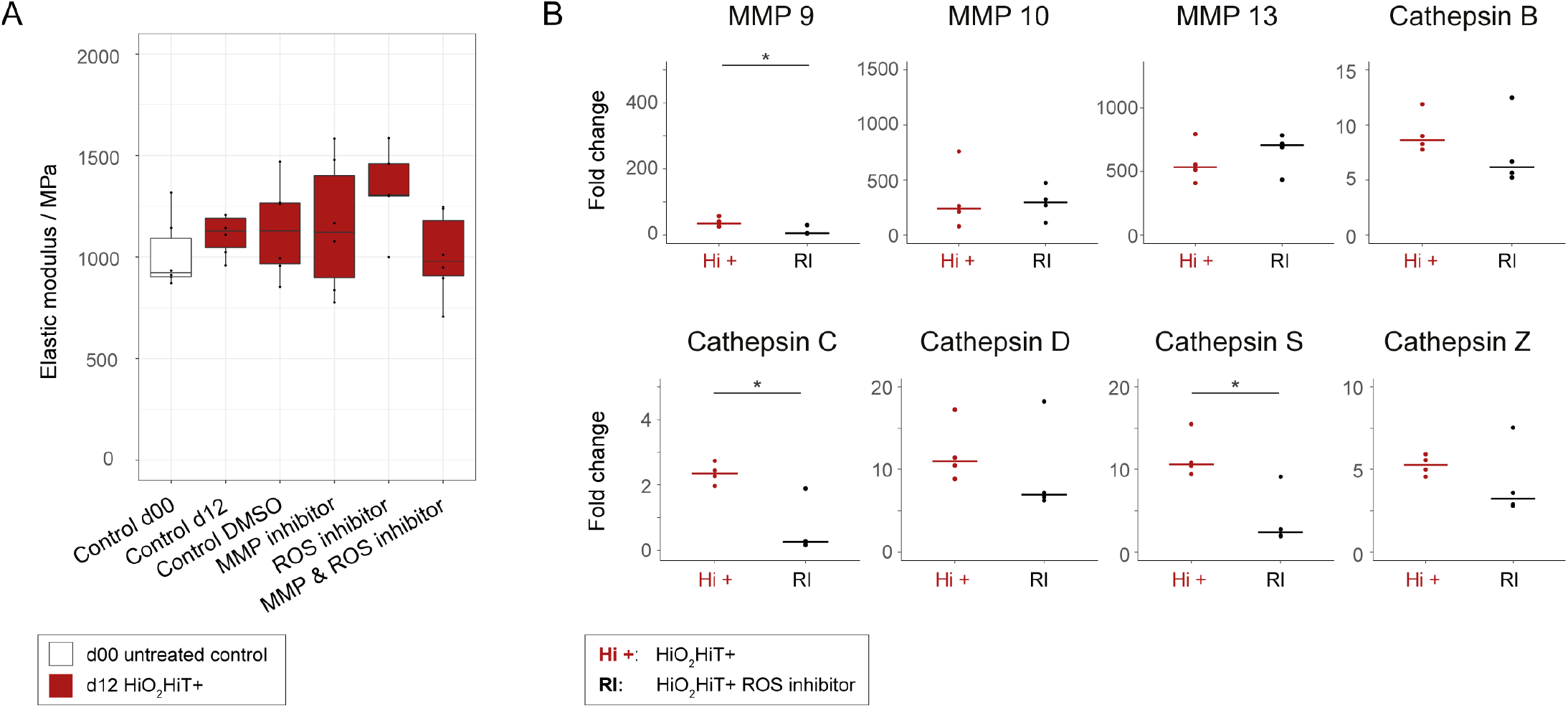
Pharmacological inhibition experiments on mechanically degrading tendon fascicles. (**A**) Elastic moduli of devitalized tissue cultured with different inhibitors at 37°C, 20kPa pO_2_ in 10% serum for 12 days. The elastic modulus is not significantly different between the conditions (n=7). Statistical analysis was performed using one-way ANOVA followed by contrast analysis in RStudio. Mean values of groups treated with inhibitory substances were compared to the control group cultured for 12 days in serum-supplemented medium (Control d12) (n = 7). (**B**) RT-PCR gene expression of proteins involved in extracellular matrix remodeling after 6 days of culture with and without the ROS inhibitor Tempol. MMP 9, cathepsins C and S are significantly down-regulated (p-value = 0.029) in tendon tissue by inhibiting ROS. Gene expression of MMP 10, 13 and cathepsins B, D and Z was not significantly affected by ROS inhibition. Displayed are the relative fold changes compared to the freshly isolated control at d00 and the horizontal crossbar represents the median. All values were normalized to the reference gene (Anxa5). Mann-Whitney U tests were performed using in RStudio by comparing the Tempol-supplemented group to the “HiO_2_HiT+” group (n = 4, p-values: * < 0.05).

## Footnotes

1 https://github.com/uzh/ezRun

2 http://www.bioinformatics.babraham.ac.uk/projects/fastqc/

3 http://www.bioinformatics.babraham.ac.uk/projects/fastq_screen/

4 https://bioconductor.org/packages/devel/bioc/manuals/DESeq2/man/DESeq2.pdf

5 https://portal.genego.com/

